# Hif-1alpha stabilisation is protective against infection in a zebrafish model of comorbidity

**DOI:** 10.1101/797480

**Authors:** Yves Schild, Abdirizak Mohamed, Edward J. Wootton, Amy Lewis, Philip M. Elks

## Abstract

Multi-drug resistant tuberculosis is a worldwide problem and there is an urgent need for host-derived therapeutic targets, circumventing emerging drug resistance. We have previously shown that hypoxia inducible-1α (Hif-1α) stabilisation helps the host to clear mycobacterial infection via neutrophil activation. However, Hif-1α stabilisation has also been implicated in chronic inflammatory diseases caused by prolonged neutrophilic inflammation. Comorbid infection and inflammation can be found together in disease settings, so it is unclear as to whether Hif-1α stabilisation would be beneficial in a holistic disease setting. Here, we set out to understand the effects of Hif-1α on neutrophil behaviour in disease-relevant settings by combining two well-characterised *in vivo* zebrafish models: TB infection (*Mycobacterium marinum* infection) and wounding (tailfin transection). We demonstrate during systemic infection, that wounding leads to increased infection burden, but the protective effect of Hif-1α stabilisation remains. A local Mm infection near to the tailfin wound site caused neutrophil migration between sites that was reduced by Hif-1α stabilisation. Our data indicate that the protective effect of Hif-1α against Mm is maintained in the presence of inflammation, highlighting its potential as a host-derived target against TB infection in a disease relevant setting.

## Introduction

Multi-drug resistance is an increasing problem worldwide and in 2017 WHO estimated that there were 490,000 cases of multi-drug resistant *Mycobacterium tuberculosis* infections (the cause of tuberculosis), alongside 600,000 new cases with resistance to the front-line drug rifampicin [1]. There is an urgent and unmet need for host-derived therapeutic targets that would circumvent the problems of emerging drug-resistance and could work in combination with current antimicrobials to completely clear patients of TB burden more rapidly [2].

Neutrophil activation is often viewed as a double-edged sword in terms of disease control [3]. Neutrophils must distinguish between sterile and infected tissue injuries to determine an appropriate response [4], one that strikes a balance between infection control and tissue damage, but the mechanisms behind this are not well understood in complex *in vivo* tissue environments, partially due to a lack of appropriate models. Damage associated molecular patterns (DAMPs) and pathogen associated molecular patterns (PAMPs) share some receptor repertoires and downstream signalling components, but there is evidence to suggest that neutrophils can differentiate between these signals [5]. Neutrophils are involved early in TB infection with influx associated with killing of bacteria in a number of cellular and animal models [3,6–8], but their function during mycobacterial infection is not well characterised. Neutrophils are important in infection control, however, they are also the drivers of many chronic inflammatory diseases such as chronic obstructive pulmonary disease (COPD) [9]. Neutrophils are one of the first immune cell types to respond to tissue injury and migrate to the wound to clear up fragments of cells and protect against pathogen invasion [10]. However, in order for wounds to heal, neutrophilic inflammation must resolve, either by programmed cell death (apoptosis), or by movement away from the wound in a process called reverse migration [11,12]. If neutrophils persist, then degranulation occurs leading to release of toxic components, further tissue damage, and consequent neutrophil recruitment; a vicious cycle of chronic inflammation that underpins many inflammatory diseases like COPD.

Chronic diseases, such as TB and COPD, often do not occur individually but exist together in patients, a situation called a comorbidity. This is especially true of TB, as one-third of the world’s population live healthily with latent TB infection for decades before a “second-hit” comorbidity leads to progression to active TB [13]. The best characterised comorbidities are co-infections with other communicable diseases, most notably HIV which causes immune deficiency and allows TB to breakout of granulomas leading to active disease [1]. However, at the same time as anti-retroviral therapy is bringing HIV under greater control, there is an alarming rise in non-communicable diseases, such as diabetes and COPD, in the same populations that have been linked to TB activation [13,14]. Many of these non-communicable diseases have an inflammatory component, yet treatment of these diseases, and indeed TB itself, is currently tailored towards the single condition rather than considering the holistic outcome of the comorbidity [15]. This is reflected in animal models, used to investigate cellular and molecular mechanisms of disease, often being based on a single condition rather than considering comorbidities, and there is a pressing need for combined models to understand the complex interactions of cells *in vivo*.

Neutrophils are exquisitely sensitive to low levels of oxygen (hypoxia), which pro-longs their lifespan and increases their bactericidal mechanisms [12,16,17]. The cellular response to hypoxia involves the activation and stabilisation of hypoxia inducible factor-1α (HIF-1α) transcription factor [18,19]. We have previously demonstrated that activating neutrophils, via stabilisation of Hif-1α, is host protective during *in vivo* mycobacterial infection; a good therapeutic outcome [20]. However, hypoxia and Hif-1α have also been shown to delay neutrophil apoptosis and reverse migration of neutrophils away from wounds in chronic inflammation models; a bad therapeutic outcome [12,21]. Therefore, the beneficial effects of Hif-1α stabilisation on a holistic-scale during infection remains unclear, due to the potential for neutrophil damage and chronic inflammation.

The zebrafish has become an invaluable animal model for TB and inflammatory disease over the last fifteen years [22]. Zebrafish embryos are transparent and development of immune transgenic lines has allowed unprecedented access to track immune cell dynamics inside an intact organism using fluorescence microscopy. Infection of zebrafish larvae with *Mycobacterium marinum* (Mm), a closely related strain to human Mtb and a natural fish pathogen, has been used to identify important molecular mechanisms involved in TB pathogenesis and granuloma formation [23]. The development of innate immune cell transgenic lines began with neutrophil labelled lines, and these have been used over the last decade in tailfin transection models to better understand the molecular mechanisms involved in both neutrophil recruitment to, and reverse migration from, a site of inflammation [12,24,25].

Here, we investigated the effects of Hif-1α stabilisation on neutrophil dynamics in dual-models of infection and wounding by combining well-characterised zebrafish Mm infection and tailfin transection models [12,20]. During systemic infection, neutrophil inflammation dynamics at the tailfin wound occur as normal while presence of a wound exacerbates infection burden. By switching to a localised infection we show that interaction between tailfin inflammation neutrophils and the site of infection occurs if cells are close enough to each other and that infection can attract neutrophils away from the tailfin wound prematurely. Stabilising Hif-1α caused preferential migration to the infection site and delayed premature neutrophil migration away from the tailfin wound to the site of infection, indicating that Hif-1α neutrophils are more sensitive to infection/wound gradients and are more likely to be retained in response to tissue challenge. Hif-1α stabilisation was effective at controlling systemic infection in the dual-model despite it prolonging neutrophil inflammation at the wound site. These data show that, on a local scale, stabilisation of Hif-1α can alter neutrophil migration dynamics, but that, on an entire organism level, the protective effect of Hif-1α stabilisation against infection remains. These findings demonstrate that comorbidities may have multiscale effects ranging from the local tissue level to the holistic level and highlight that the zebrafish is a promising model to investigate both levels of effects. Although stabilisation of Hif-1α has detrimental effects on neutrophil inflammation resolution, the dual-model highlights that it is a promising drug target against TB, even in the presence of an inflammatory comorbidity.

## Materials and methods

### Zebrafish husbandry

All the zebrafish used in this project were raised in the University of Sheffield Home Office approved aquarium and were kept under standard protocols as previously outlined [26]. Adult zebrafish were kept in tanks of no more than 40 adult fish, and experience a 14-hour light and 10-hour dark cycle. A recirculating water supply is maintained and the temperature of the water is kept at 28°C. Embryos for this study were generated by in-crossing *TgBAC(mpx:Gal4.VP16);Tg(UAS:Kaede)i222* or *Tg(mpx:GFP)i114* [25,27].

### Ethics

All procedures over the course of this project were performed on embryos that were less than 5.2 days post fertilisation (dpf) and were therefore considered outside of the Animals (Scientific Procedures) Act. Procedures were carried out to standards set by the UK Home Office on the Project Licence P1A4A7A5E held by Professor Stephen Renshaw at the University of Sheffield.

### Tailfin transection

For all experiments, larval tailfins were transected at 48 hours post fertilisation (hpf) as previously described [12]. Kaede-expressing wound neutrophils were photoconverted at 4 hours post wound (hpw) using a SOLA light engine white light LED (Lumencor, USA) through DAPI filters on a Leica DMi8 inverted widefield microscope (Leica Microsystems, Germany). Timelapse microscopy was performed using a Leica DMi8 inverted widefield microscope (Leica Microsystems, Germany) using a HC FL PLAB 10x/0.40 lens and captured using a Hammamatsu ORCA-Flash 4.0 camera (Hammamatsu, Japan). Neutrophil counts were performed with the investigator blinded to the experimental group on a Leica MZ10 F Stereomicroscope with fluorescence (Leica Microsystems, Germany).

### *Mycobacterium marinum* infection

Mm infection experiments were performed using *M. marinum* M (ATCC #BAA-535), containing a psMT3-mCherry or psMT3 mCrimson vector [28]. Injection inoculum was prepared from an overnight liquid culture in the log-phase of growth resuspended in 2% polyvinylpyrrolidone40 (PVP40) solution (CalBiochem) as previously described [20].

For systemic infection 150-200 colony forming units (CFU) were injected into the caudal vein at 28-30hpf, as previously described [29].

For localised somite infection, fish were anaesthetised in 0.168 mg/ml Tricaine (Sigma-Aldrich) and were microinjected with 500CFU (colony forming units) of Mm in the 26^th^-27^th^ somite [30].

### Hif-1α stabilisation

Embryos were injected with dominant active *hif-1αb* (ZFIN: hif1ab) variant RNA at the one cell stage as previously described [12,31]. Phenol red (PR) (Sigma Aldrich) was used as a vehicle control.

Hif-1α was stabilised pharmacologically using hydroxylase inhibitors FG4592, 5μM or DMOG, 100μM (dimethyloxaloylglycine), with DMSO control.

### Bacterial pixel count

Infected zebrafish larvae were imaged at 4 days post infection (dpi) on an inverted Leica DMi8 with a 2.5× objective lens. Brightfield and fluorescent images were captured using a Hammamatsu OrcaV4 camera. Bacterial burden was assessed using dedicated pixel counting software as previously described [20,32].

### Image and Statistical Analysis

Microscopy data was analysed using Leica LASX (Leica Microsystems, Germany) and Image J software. All data were analysed (Prism 7.0, GraphPad Software) using t-tests for comparisons between two groups and one-way ANOVA (with Bonferonni post-test adjustment) for other data. P values shown are: \**P* < .05, \*\**P* < .01, and \*\*\**P* < .001.

## Results

### Infection induced neutrophil emergency haematopoeisis and increased neutrophilic inflammation to the detriment of infection control

Infection and inflammation commonly occur in the same individual during disease, yet many *in vivo* experimental systems investigate immune responses to these processes independently of each other. We set out to develop *in vivo* zebrafish models of infection and inflammation, that we have termed “dual-models”. Initially we combined two well-defined models; a *Mycobacterium marinum* (Mm) model of systemic infection (injection of bacteria into the caudal vein at 30-32 hours post fertilisation (hpf) and assessing bacterial burden at 4 days post infection (dpi)) and a tailfin wound model of neutrophilic inflammation (transection of the tailfin at 2 days post fertilisation (dpf) with neutrophil inflammation resolving at 24 hours post wound (hpw)) [33,34] (Figure 1A). We first assessed whether injury at the caudal vein (the site of Mm infection) caused by the microinjection process itself would affect neutrophil behaviour at the tailfin wound. Injection of PVP into the caudal vein (mock infection control) caused no difference to the number of neutrophils at the peak of recruitment to the tailfin wound (6hpw), nor after neutrophil inflammation resolution at 24hpw (not injected, NI, compared to PVP injected) (Figure 1B).

**Figure 1:**
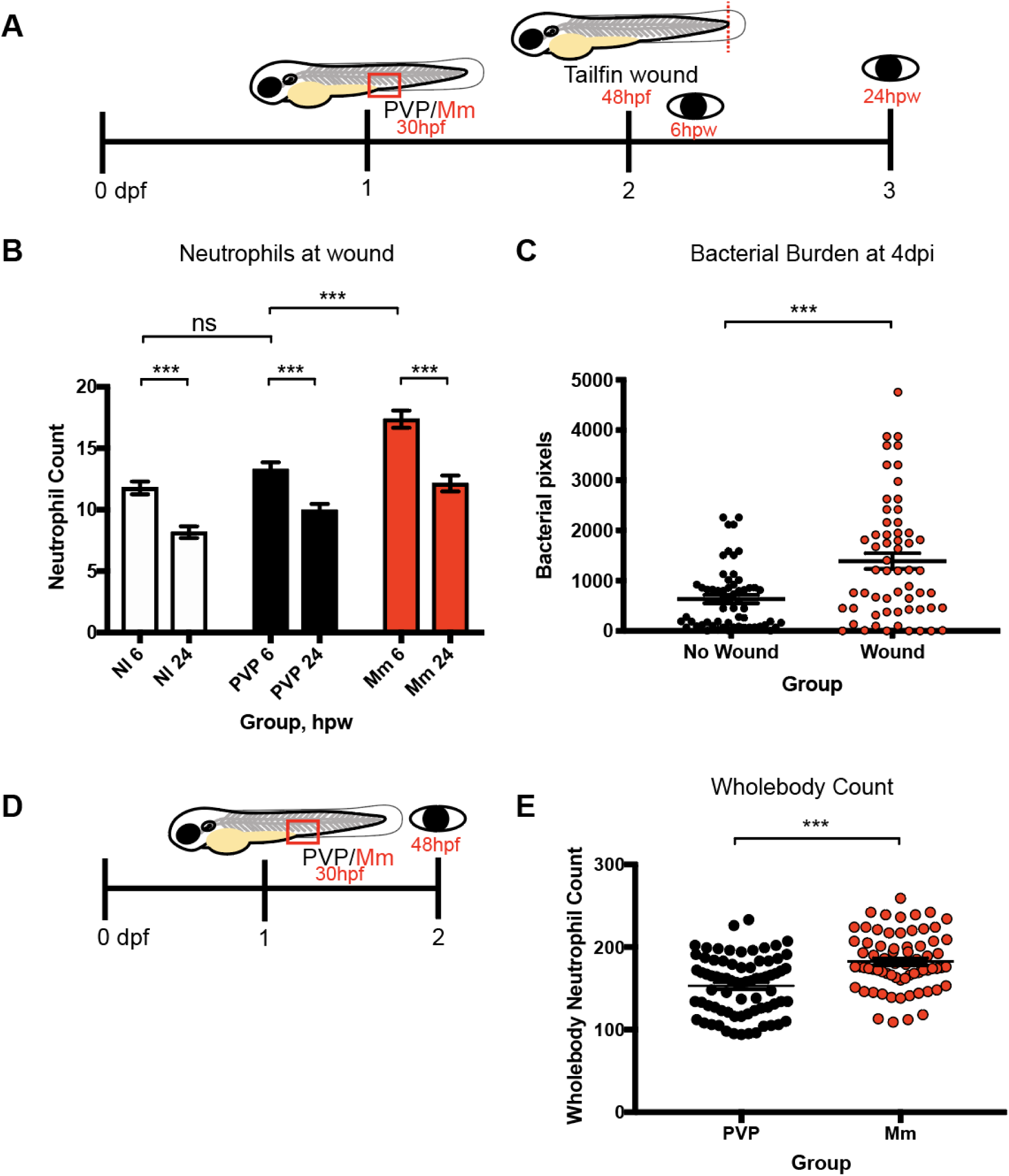
Mm Infection induced neutrophil emergency haematopoeisis and increased neutrophilic inflammation to the detriment of infection control. (A) Schematic of experiment for B-C. (B) Neutrophil numbers at the wound at 6 and 24 hours post wound (hpw). Groups are not injected (NI), control injection with PVP (PVP) and Mm injection (Mm). Data shown are mean ± SEM, n=75-85 accumulated from 3 independent experiments. (C) Bacterial burden of larvae with or without a tailfin wound. Data shown are mean ± SEM, n=58 accumulated from 3 independent experiments. (D) Schematic of experiment for E. (E) Total, whole-body neutrophil numbers at 2dpf, after 18 hours post infection (hpi) with PVP or Mm. Data shown are mean ± SEM, n=69-74 accumulated from 3 independent experiments.

The presence of systemic Mm infection increased neutrophil number at the wound at both the 6hpw and 24hpw timepoints compared to NI and PVP controls (Figure 1B). Although overall neutrophil numbers were increased by infection at 6hpw and 24hpw, the resolution of neutrophil inflammation still occurred (Figure 1B). Infection levels were measured in the dual-model using fluorescent Mm and assessing bacterial burden at 4dpi. Levels of Mm infection were significantly increased in the presence of neutrophilic inflammation at the wound site compared to non-wounded controls (Figure 1C) indicating that the presence of localised tailfin inflammation is detrimental to infection control. We assessed whole body neutrophil counts after Mm infection without a tailfin injury and confirmed that total neutrophil number was increased after Mm infection (Figure 1D-E) consistent with emergency haematopoeisis [35].

### Neutrophils distributed to local infection and wound sites

To investigate neutrophil migration to infection and wound stimuli in a dual-model we challenged 3dpf zebrafish larvae with a tailfin wound immediately followed by a local somite infection into the 26-27^th^ somite (Mm or PVP mock infection control) and counted neutrophils at each site over time (Figure 2A). When challenged with Mm infection alone or tailfin wound alone, neutrophils from the caudal haematopoietic tissue (CHT) and surrounding areas migrated to each respective site and peaked at 4-6hpw/i (Figure 2B-D). Of note, some neutrophils were present at the site of infection before challenge (on average 10 neutrophils) due to the natural distribution of neutrophils at this stage, with very few present at the end of the tail (the wound site, <5 neutrophils) (Figure 2B-D). When tailfin wounding was followed by PVP injection (as a mock infection control), neutrophils migrated to both the somite PVP site and the tailfin wound site, indicating that a wound in the somite was sufficient to attract neutrophils, while neutrophils were still able to migrate beyond this to the tailfin wound (although to a lesser extent than wound alone, Figure 2B-D). When tailfin wound was followed by somite Mm infection, neutrophils migrated to the somite infection site at the expense of tailfin wound neutrophils (Figure 2B-D). These data indicate that the signal gradient caused by Mm infection is additive to that of the somite injury alone and that neutrophils preferentially migrate to Mm and are retained at infection rather than travelling further along the trunk to the tailfin wound.

**Figure 2:**
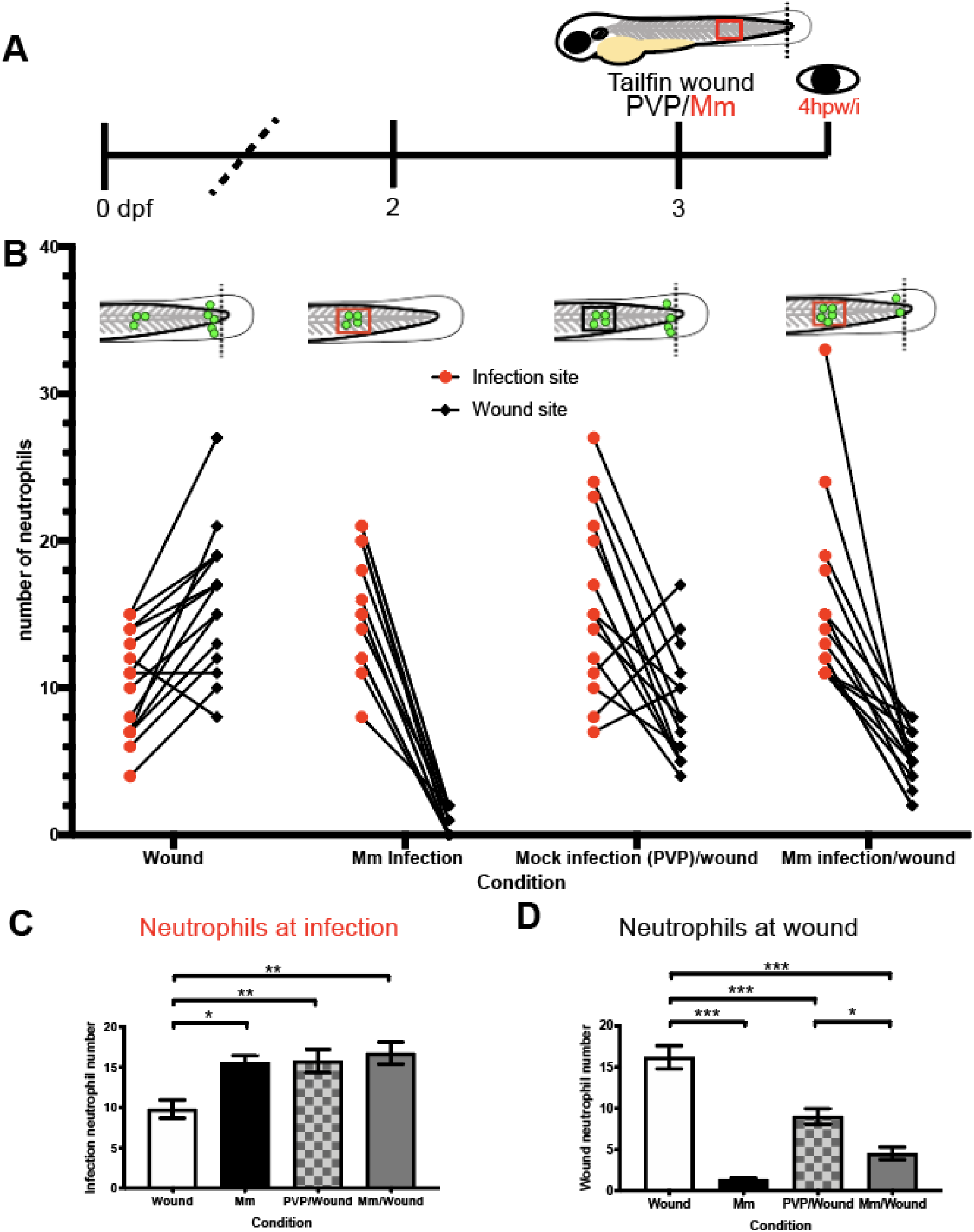
Neutrophils distributed to local infection and wound sites. (A) Schematic of experiment for B-D. (B) Number of neutrophils at site of infection and tailfin wound at 4hpi/w. Data shown are mean ± SEM, n=9-13 representative of 3 independent experiments. (C) Neutrophil numbers at the infection site at 4hpi. Data shown are mean ± SEM, n=9-13 representative of 3 independent experiments. (D) Neutrophil numbers at the wound site at 4hpi. Data shown are mean ± SEM, n=9-13 representative of 3 independent experiments.

### Neutrophils preferentially migrated to a new infection stimulus rather than patrol a wound site

In a single model of tailfin wound, once neutrophils have migrated to a wound site (between 1-6hpw), they are retained at the wound, patrolling until the resolution phase of inflammation (6-12hpw) [12,25]. We have previously demonstrated that neutrophils migrate away from the wound by a diffusion process at around 8-12hpw when neutrophils become desensitised to signals that retains them at the wound [36]. We hypothesised that infection can overcome this retention signal at the wound site and attract neutrophils prematurely away from the wound. We therefore developed a dual model where, at 4hpw, a localised Mm infection was introduced into the 26-27^th^ somite (Figure 3A). 4hpw is a timepoint at which neutrophils are still being recruited to the wound and would not have started to reverse migrate away in a single wound model, a process that normally occurs after 6-12hpw [10,12]. Photoconversion of *Tg(mpx:Gal4/UAS:Kaede)* neutrophils at the tailfin wound at 4hpw allowed identification of neutrophils that had visited the wound (“wound experienced” red neutrophils), compared to those that had not (“wound naïve” green neutrophils) (Figure 3B). We demonstrated that injection of Mm into the 26-27th somite was sufficient to attract neutrophils away from the wound (wound experienced neutrophils) between 4hpw-6hpw (Figure S1). By 100mpc (minutes post conversion) almost all wound-experienced neutrophils had been attracted away from the tailfin wound by infection (Figure 3D). These data demonstrate that the “second hit” of infection was sufficient to overcome signalling that retains neutrophils at the initial tailfin wound site.

**Figure 3:**
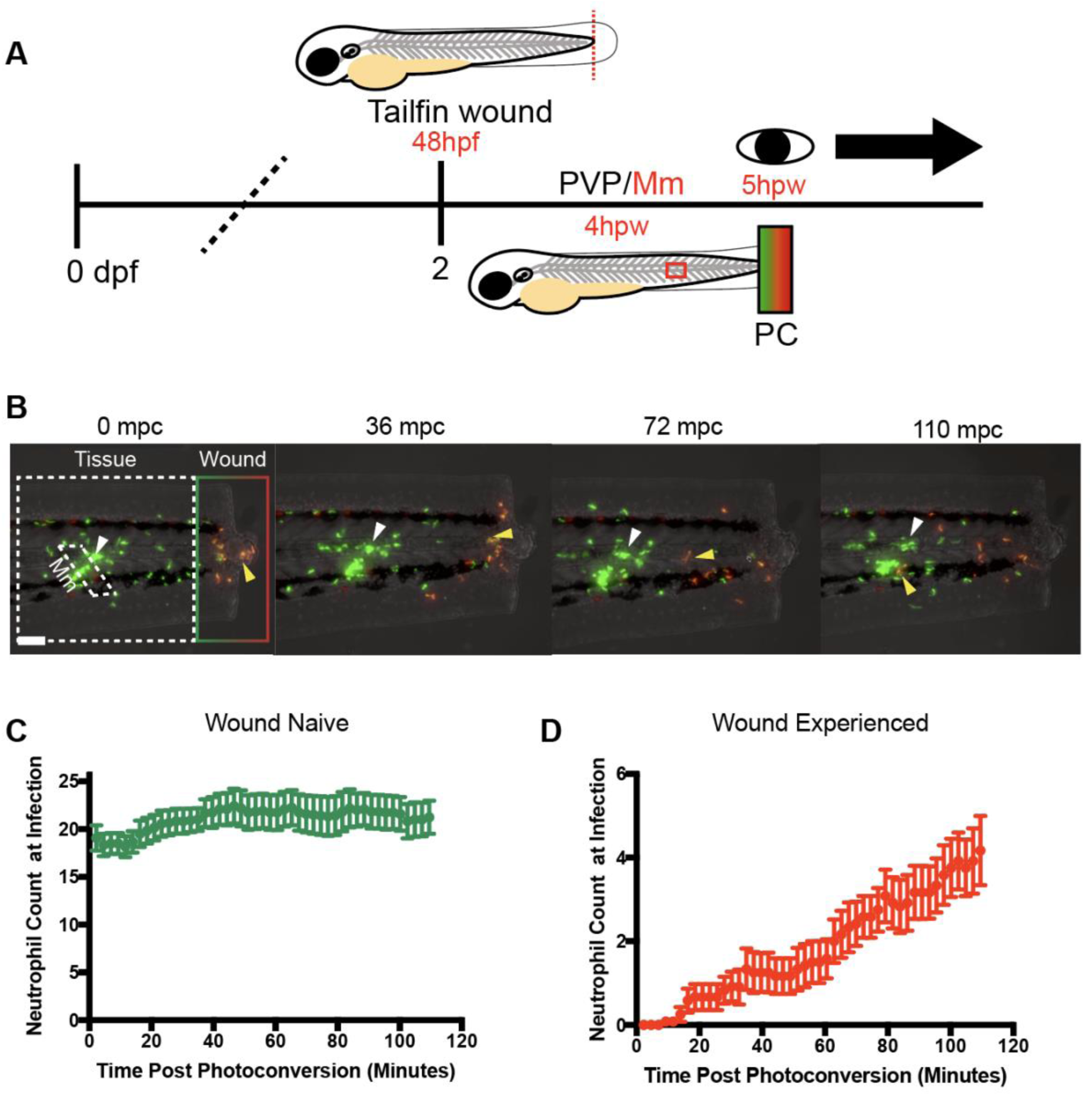
Neutrophils preferentially migrated to a new infection stimulus rather than patrol a wound site. (A) Schematic of experiment for B-D. (B) Stereo-fluorescence micrographs of a tailfin transected embryo after 26/27^th^ somite infection with Mm. Wound-naïve neutrophils are green only and those photoconverted at the wound at timepoint zero (wound-experienced) begin as red-only and regain GFP (therefore giving a yellow overlay) over the course of the timelapse as nascent Kaede fluorescent protein is made. Both wound-naïve (white arrowhead) and wound-experienced (yellow arrowhead) are recruited to the localised site of Mm infection before 110 minutes post conversion (mpc), even though the timelapse is begun at 5hpw, a timepoint when neutrophils would normally still be recruited to the tailfin transection. (C) Number of green, wound-naïve neutrophils at infection site over 1.5hpi. Data shown are mean ± SEM, n=12 embryos accumulated from 3 independent experiments. (D) Number of red, wound-experienced neutrophils at infection site over 1.5hpi. Data shown are mean ± SEM, n=12 embryos accumulated from 3 independent experiments.

### Hif-1α stabilisation retained neutrophils at infection at the expense of migration to tailfin wound

Hypoxia signalling, via stabilisation of Hif-1α, has profound effects on neutrophil behaviours and antimicrobial activity [12,20,21]. We set out to understand whether Hif-1α stabilisation affected neutrophil behaviour in our dual models of infection and inflammation. Endogenous Hif-1α was stabilised pharmacologically using the hydroxylase inhibitors FG4592 and DMOG [12] 4 hours before infection with Mm into the 26-27th muscle somite. This was followed by immediate tailfin wound and neutrophil numbers were counted at each site at 6pw/I (Figure 4A). The solvent control for both hydroxylase inhibitors (DMSO), caused no difference in neutrophil migration to infection and wound at 6hpw/i compared to untreated larvae (Figure 4B-F). Treatment with either FG5492 or DMOG caused significantly increased neutrophil migration to the infection site with fewer neutrophils migrating to the tailfin wound compared to DMSO controls (Figure 4B-F). These findings were confirmed by genetic stabilisation of Hif-1α using dominant active Hif-1α (Figure 4G-I). These data suggest that neutrophils primed with Hif-1α are more sensitive to the local infection chemokine gradient at the expense of the more distant gradient emanating from the wound.

**Figure 4:**
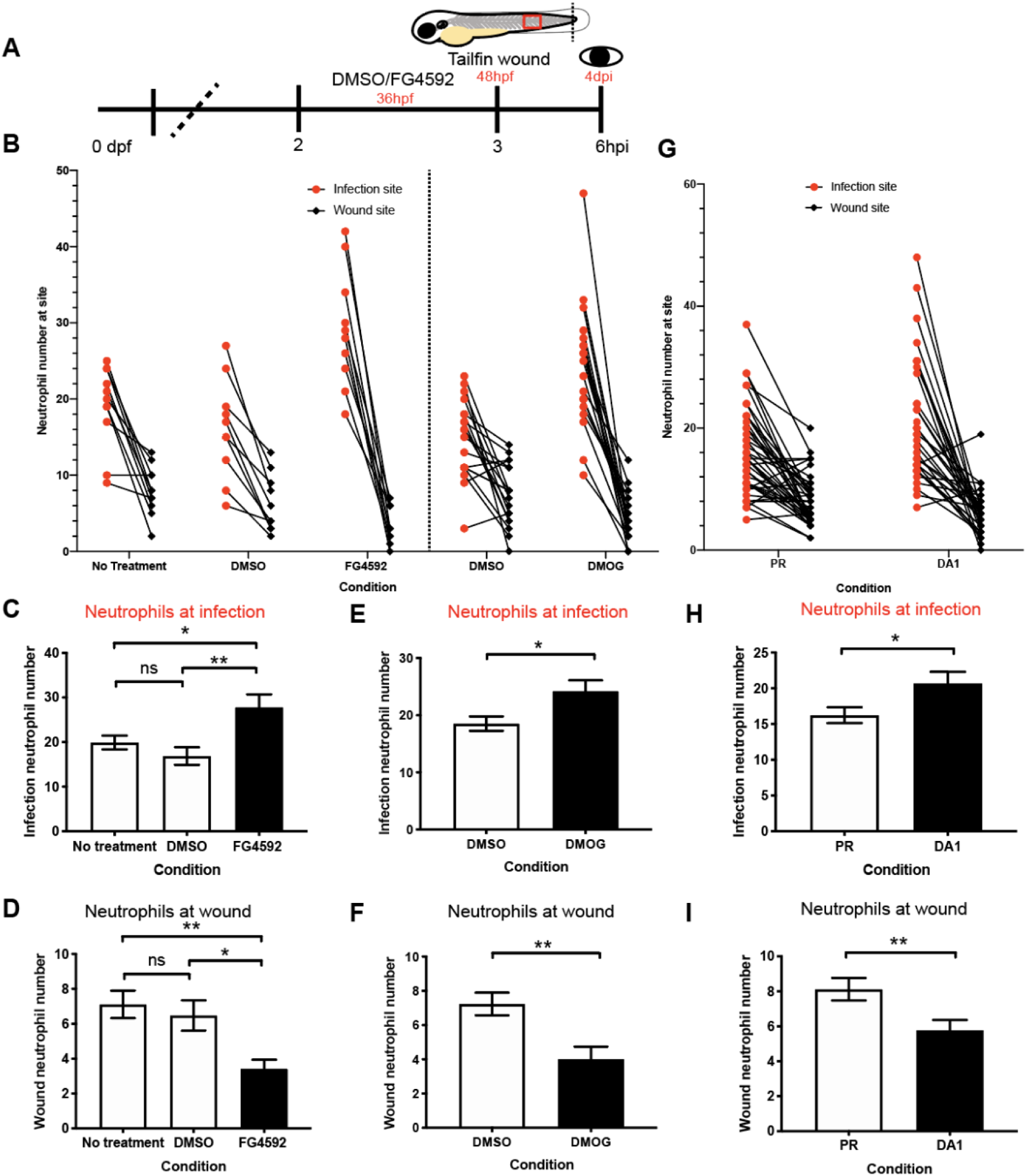
Hif-1α stabilisation retained neutrophils at infection at the expense of migration to tailfin wound. (A) Schematic of experiment for B-F. (B) Number of neutrophils at site of infection and tailfin wound at 4hpi/w after Hif-1α stabilisation with FG4592 or DMOG with No treatment and DMSO controls. Data shown are mean ± SEM, n=9-15 representative of 3 independent experiments. (C) Neutrophil numbers at the infection site at 4hpi with DMSO and FG4592 treatment. Data shown are mean ± SEM, n=10-11 representative of 3 independent experiments. (D) Neutrophil numbers at the wound site at 4hpi with DMSO and FG4592 treatment. Data shown are mean ± SEM, n=10-11 representative of 3 independent experiments. (E) Neutrophil numbers at the infection site at 4hpi with DMSO and DMOG treatment. Data shown are mean ± SEM, n=19 representative of 3 independent experiments. (F) Neutrophil numbers at the wound site at 4hpi with DMSO and DMOG treatment. Data shown are mean ± SEM, n=19 representative of 3 independent experiments. (G) Number of neutrophils at site of infection and tailfin wound at 4hpi/w after Hif-1α stabilisation with dominant active Hif-1α (DA1) or phenol red (PR) controls. Data shown are mean ± SEM, n=20-22 representative of 3 independent experiments. (H) Neutrophil numbers at the infection site at 4hpi with PR and DA1. Data shown are mean ± SEM, n=19 representative of 3 independent experiments. (I) Neutrophil numbers at the wound site at 4hpi with PR and DA1 treatment. Data shown are mean ± SEM, n=36-41 accumulated from 3 independent experiments.

### Hif-1α stabilisation delayed wound-experienced neutrophil migration to Mm infection

We have previously demonstrated, in a single tailfin wound model, that stabilisation of Hif-1α delays neutrophil reverse migration away from the wound [12]. However, here we show that a local Mm infection is able to attract neutrophils away from the tailfin wound prematurely (Figure 3). We therefore hypothesised that Hif-1α would prevent wound-experienced neutrophils from exiting the injury site prematurely to migrate to a localised infection site. Wound-naïve neutrophil attraction to the site of Mm infection was not altered by DA Hif-1α compared to phenol red (PR) controls (Figure 5A-B). Infection was sufficient to attract wound-experienced neutrophils away from the wound prematurely, but DA Hif-1α neutrophils were significantly delayed in their migration towards localised Mm infection compared to PR controls (Figure 5B-C). The migration speed of wound-experienced neutrophils was lower in the DA Hif-1α group compared to the PR group, largely due to their tighter association to the wound edge and less migration away (Figure 5D). This decrease in migration speed was more marked in wound-experienced neutrophils that were successful in migrating away from the wound edge towards the Mm infection site (Figure 5E). These neutrophils migrated to the infection site at two-thirds of the speed in DA Hif-1α embryos compared to the PR controls (Figure 5E). Furthermore, they took a less direct route to the infection, with the meandering index of these neutrophils significantly lower in the DA Hif-1α group compared to PR controls (Figure 5F). These data indicate that Hif-1α stabilised neutrophils remain more sensitive to the wound signalling gradient, even if successful in escaping the wound to a second hit of infection. It is interesting to note that, in many cases, wound-experienced neutrophils migrating away from the wound in the DA Hif-1α group dithered between the wound and infection sites, with a shuttling movement backwards and forwards, a behaviour not observed in PR controls (Movie S1). Dithering between infection and wound sites was also not observed in DA Hif-1α wound-naïve neutrophils in the same individual larvae, suggesting a difference between wound-experienced and wound-naïve neutrophils in their detection of the two stimuli.

**Figure 5:**
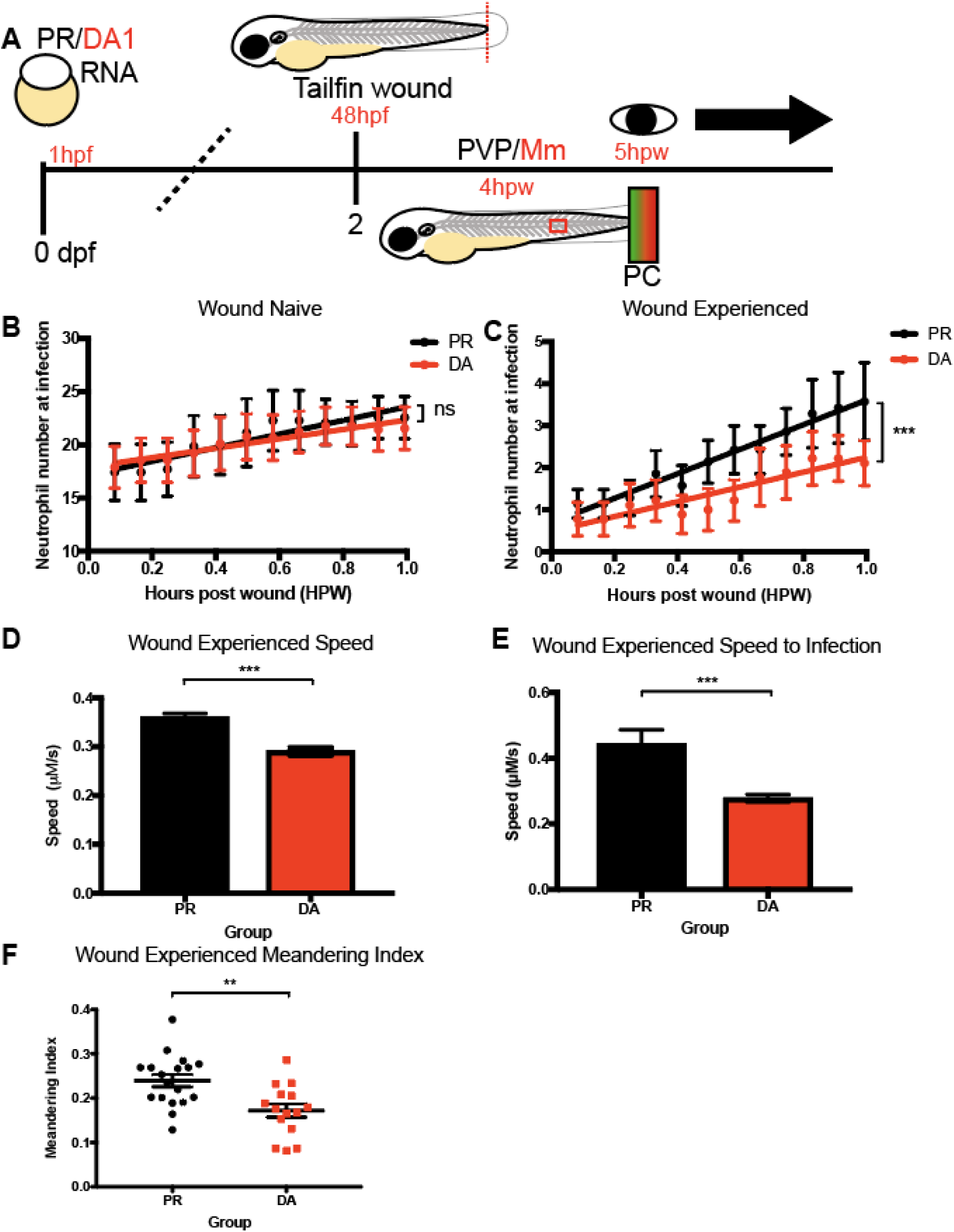
Stabilisation of Hif-1α delayed the migration of wound-experienced neutrophils to a local site of Mm infection. (A) Schematic of experiment for B-F. (B) Number of green, wound-naïve neutrophils at infection site over 1 hour post wound (hpw). Groups shown are DA Hif-1α (DA, red points) and phenol red controls (PR, black points). Data shown are mean ± SEM, n=7-9 embryos accumulated from 3 independent experiments. Line of best fit shown is calculated by linear regression. P value shown is for the difference between the 2 slopes. (C) Number of red, wound-experienced neutrophils at infection site over 1 hpw. Groups shown are DA Hif-1α (DA, red points) and phenol red controls (PR, black points). Data shown are mean ± SEM, n=7-9 embryos accumulated from 3 independent experiments. Line of best fit shown is calculated by linear regression. P value shown is for the difference between the 2 slopes. (D) Speed of red, wound-experienced neutrophil movement at the wound site. Groups shown are DA Hif-1α (DA) and phenol red controls (PR). Data shown are mean ± SEM, n=5-6 embryos accumulated from 3 independent experiments. (E) Speed of red, wound-experienced neutrophils migrating from the wound site to the infection site. Groups shown are DA Hif-1α (DA) and phenol red controls (PR). Data shown are mean ± SEM, n=5-6 embryos accumulated from 3 independent experiments. (F) Meandering index of red, wound-experienced neutrophils migrating from the wound site to the infection site. Groups shown are DA Hif-1α (DA) and phenol red controls (PR). Data shown are mean ± SEM, n=15-18 embryos accumulated from 2 independent experiments.

Taken together, these data indicate that wound-experienced neutrophils in Hif-1α stabilised larvae remain more sensitive to the wound gradient and are less likely to migrate to the second hit infection site compared to normal controls.

### Mm burden was decreased by Hif-1α stabilisation, despite delayed resolution of neutrophilic inflammation

In the single model of Mm infection we have previously shown that Hif-1α stabilisation reduced bacterial burden; a good therapeutic outcome [20]. However, in the single tailfin model, Hif-1α delayed neutrophil inflammation resolution away from the wound; a bad therapeutic outcome in diseases of chronic inflammation [12]. As infection and chronic inflammation are common attributes of comorbidities, we investigated whether the beneficial therapeutic outcome of Hif-1α stabilisation in infection would be maintained in the presence of chronic inflammation.

We observed an increase in neutrophil recruitment to the tailfin wound after Mm infection (at 6hpw) in PR controls (Figure 6A-B), in keeping with the emergency hematopoietic effect of infection observed earlier (Figure 1E). No effect of DA Hif-1α was observed on neutrophil recruitment compared to PR controls (Figure 6B), consistent with previous observations in the single tailfin transection model [12]. Neutrophil numbers at the wound after resolution, at 24hpw were increased by DA Hif-1α compared to PR controls in the presence (Mm) or absence (PVP) of Mm infection (Figure 6C) and the percentage resolution (6-24hpw) was reduced by Hif-1α stabilisation compared to PR controls (Figure 6D), indicating that Hif-1α stabilisation delays neutrophil inflammation resolution in the presence of systemic infection.

**Figure 6:**
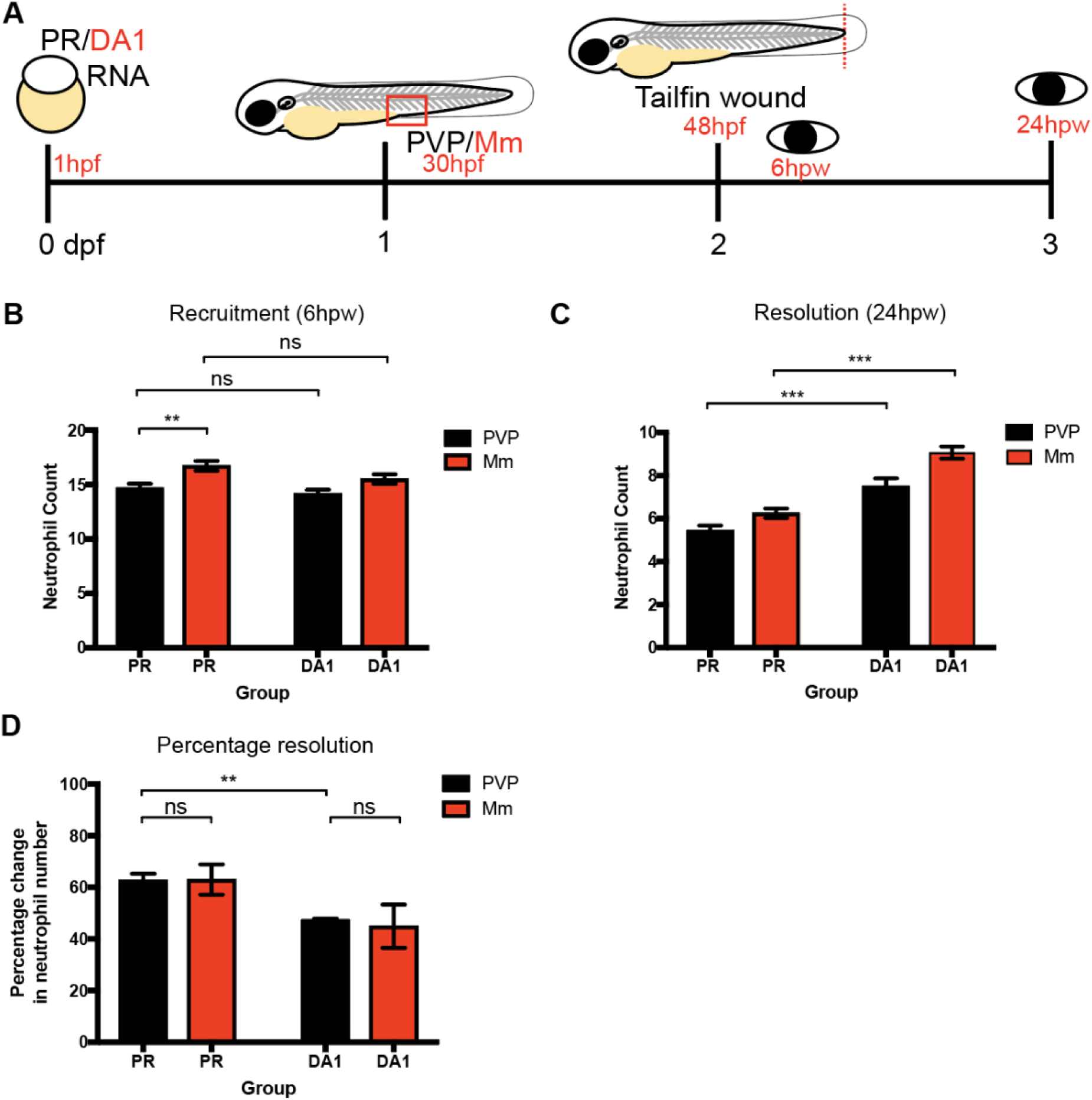
Stabilisation of Hif-1α delayed neutrophil inflammation resolution in the presence of systemic Mm infection. (A) Schematic of experiment for B-D. (B) Neutrophil numbers recruited to the tailfin wound at 6hpw. Groups are phenol red (PR) and DA Hif-1α (DA) injected at 30hpf with PVP or Mm. Data shown are mean ± SEM, n=62-111 accumulated from 3 independent experiments. (C) Neutrophil numbers at the tailfin wound at 24hpw. Groups are phenol red (PR) and DA Hif-1α (DA) injected at 30hpf with PVP or Mm. Data shown are mean ± SEM, n=62-111 accumulated from 3 independent experiments. (D) Percentage resolution of neutrophil inflammation (between 6 to 24 hpw). Groups are phenol red (PR) and DA Hif-1α (DA) injected at 30hpf with PVP or Mm. Data shown are mean ± SEM, n=62-111 accumulated from 3 independent experiments.

DA Hif-1α larvae had decreased bacterial burden compared to PR controls indicating that the protective effects of Hif-1α stabilisation remained, even in the presence of tailfin inflammation (Figure 7A-C). This is despite our finding that an inflammatory process (tailfin wound) during systemic infection caused a marked increase in infection levels in the absence of Hif-1α stabilisation (Figure 7B-C). These results indicate that Hif-1α remains protective against Mm even when neutrophil inflammation resolution is delayed at the tailfin.

**Figure 7:**
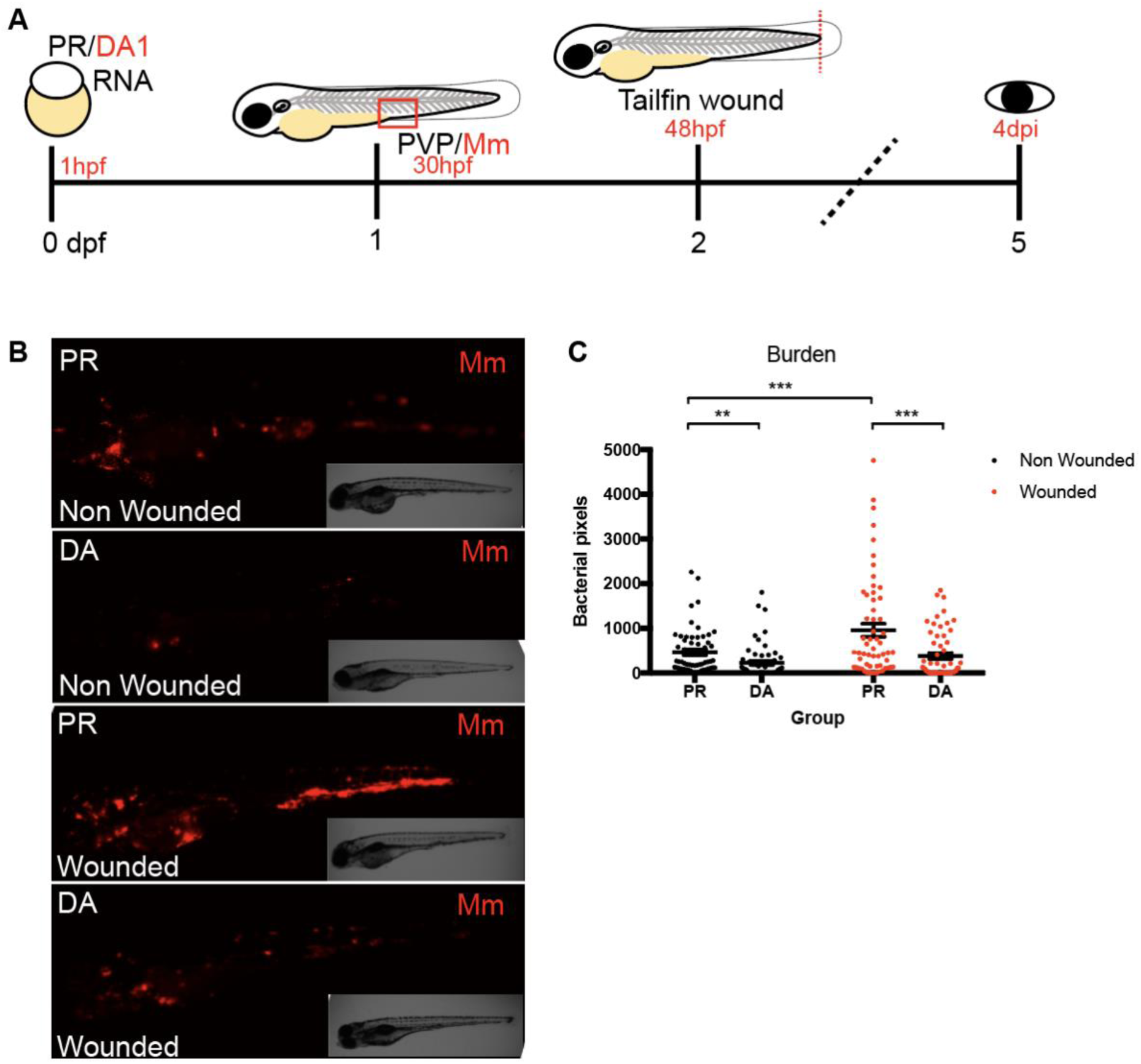
Mm burden was increased after tailfin wounding but control by Hif-1α stabilisation was maintained. (A) Schematic of experiment for B-C. (B) Stereo-fluorescence micrographs of Mm mCherry infected 4dpi larvae after injection with DA Hif-1α (DA1) and phenol red (PR) as a negative control and either wounded at 48hpf or non-wounded. (C) Bacterial burden of larvae shown in (B). Data shown are mean ± SEM, n=58 accumulated from 3 independent experiments.

## Discussion

With the emergence of antibiotic resistance, there is increasing interest to find host-derived factors that could act as therapeutic targets [2]. We have previously identified targeting neutrophils in zebrafish *in vivo* models of tuberculosis infection as a mechanism to decrease infection burden via Hif-1α stabilisation [20]. Physiological hypoxia and Hif-1α stabilisation have been demonstrated to have activating effects on neutrophils in a growing number of models, increasing their antimicrobial capabilities *in vitro, ex vivo* and *in vivo* [16,17]. These findings have been tempered by clinical observations that activated neutrophils are associated with chronic disease, leading to excess tissue damage and poor disease outcomes [11,21]. Signs of Hif-1α stabilisation being detrimental to inflammation resolution were also observed in a zebrafish tailfin wound where resolution of neutrophil inflammation is delayed, however no further adverse defects were seen [12]. Patient studies address neutrophil behaviour at the chronic stages of disease by which time there is a cycle of neutrophil overactivation, degranulation, tissue damage and further recruitment. Targeting neutrophils at earlier disease stages could therefore be highly beneficial before this chronic cycle can begin, but effects in patients with comorbid TB with inflammatory conditions such as COPD are unclear. Here we address the roles of activated neutrophils at infection and wound sites in an individual organism as a model of comorbid infection and inflammation.

We developed dual-infection/inflammation models to investigate the effects of Hif-1α on neutrophil migration to wound and infection sites simultaneously. Using localised Mm infection and tailfin wound we found that neutrophils dispersed between infection and wound sites, but that when Hif-1α was stabilised, neutrophils seldom migrated past the local infection to the tailfin wound. Hif-1α stabilisation also retained neutrophils at the tailfin wound when a second hit of infection was introduced, while in wildtype larvae infection caused premature migration away from the wound to the infection site. These data indicate that Hif-1α stabilisation causes increased sensitivity to wound or infection gradients, leading to retention of neutrophils and reduced ability of these cells to respond to competing signals.

Wound-naïve neutrophils were able to migrate to Mm at the same rate when Hif-1α is stabilised, while wound-experienced neutrophils are slower to respond and remain at the wound for longer. In some instances, when Hif-1α is stabilised the neutrophils seem unable to decide which stimuli to migrate to, shuttling between the two sites. Hif-1α stabilisation caused no effect on neutrophil recruitment to the tailfin wound in the single inflammation model, therefore is unlikely to have effects on recruitment signalling [12]. Taken together, these data indicate that recognition of “retention signals” by neutrophils is sensitised by stabilised Hif-1α, keeping neutrophils at the wound or infection site, and that there is an as yet unidentified molecular change in Hif-1α stabilised neutrophils that alters their sensitivity to these tissue gradients. Likely candidates for Hif-1α targets include G protein coupled receptors (GPCRs) that are involved in neutrophil migration (many chemokine receptors are GPCRs) and are regulated by Hif-1α in immune cells (eg., CXCR1, CXCR2 or CXCR4) [37–41]. Cxcr1/2 have been implicated in retention of neutrophils at a tailfin wound in zebrafish and we have recently demonstrated that decreasing Cxcr4 signalling causes premature reverse migration away from the tailfin wound [42].

We combined well-characterised models to address the outcomes of Hif-1α stabilisation on infection and inflammation. As well as demonstrating that the protective effect of Hif-1α stabilisation during infection being maintained with a wound present, it is interesting to note that a tailfin wound was deleterious to the host, increasing the burden of Mm infection. We have previously demonstrated, in single models of wounding that there is robust upregulation of pro-inflammatory Il-1β in neutrophils after both Hif-1α stabilisation and wounding [43,44]. These data indicate that stimulation of neutrophils by wounding and Hif-1α have differential effects on the outcome of infection, and that if neutrophils are appropriately activated it can be beneficial on a whole-organism scale.

Previous work from our group demonstrated that, during the reverse migration phase, (>12hpw) wound-experienced neutrophils reverse migrating away from the wound towards a range of infection stimuli (*Staphylococcus aureus* and zymosan) display unaltered migration behaviour compared to nearest-neighbour, wound-naive neutrophils [30]. In the absence of Hif-1α stabilisation, this appears to be the case in our Mm/wounding model, with both wound-naïve and wound-experienced neutrophils able to respond to the secondary local infection. However, when Hif-1α is stabilised differences in neutrophil migration behaviour become evident, and wound-experienced neutrophils change behaviour and are slower to migrate to the second hit, while wound-naïve neutrophils migrate as normal, indicating that neutrophils that have visited the wound can differ from those that have not.

We kept as many aspects of each individual model as close as possible to those published previously in order to avoid setting up a dual-model with undefined individual characteristics that would potentially complicate interpretation [12,20]. As investigations of comorbidities increase we anticipate that dual-models will increase in popularity, but with a plethora of possible combinations and timings of stimuli available, care will be required to understand the relevance of these models to disease situations.

Using dual-models of infection and wounding we have highlighted that comorbidity is likely to have a range of effects on neutrophil behaviour during infection that differ on the local tissue scale compared to the whole-organism, holistic level. Although Hif-1α stabilisation could be detrimental at local level inflammation, our dual-models suggest that on a whole-organism level neutrophil activation by α stabilisation is not harmful and could be a promising host-derived treatment strategy against TB.

## Acknowledgements

The authors would like to thank The Bateson Aquarium Team for fish care and the IICD Technical Team for practical assistance (University of Sheffield). Thanks to Stephen Renshaw (University of Sheffield) for constructive comments on the manuscript.

## Author Contributions

Conceived and designed the experiments: YS, AM, EJW, PME. Performed the experiments: YS, AM, EJW, AL, PME. Analyzed the data: YS, AM, EJW, PME. Wrote the paper: PME.

