## Supplementary Figures and legends for "Hif-1alpha stabilisation is protective against infection in a zebrafish model of comorbidity"

**Supplemental Figures and Movie**

### Figure S1

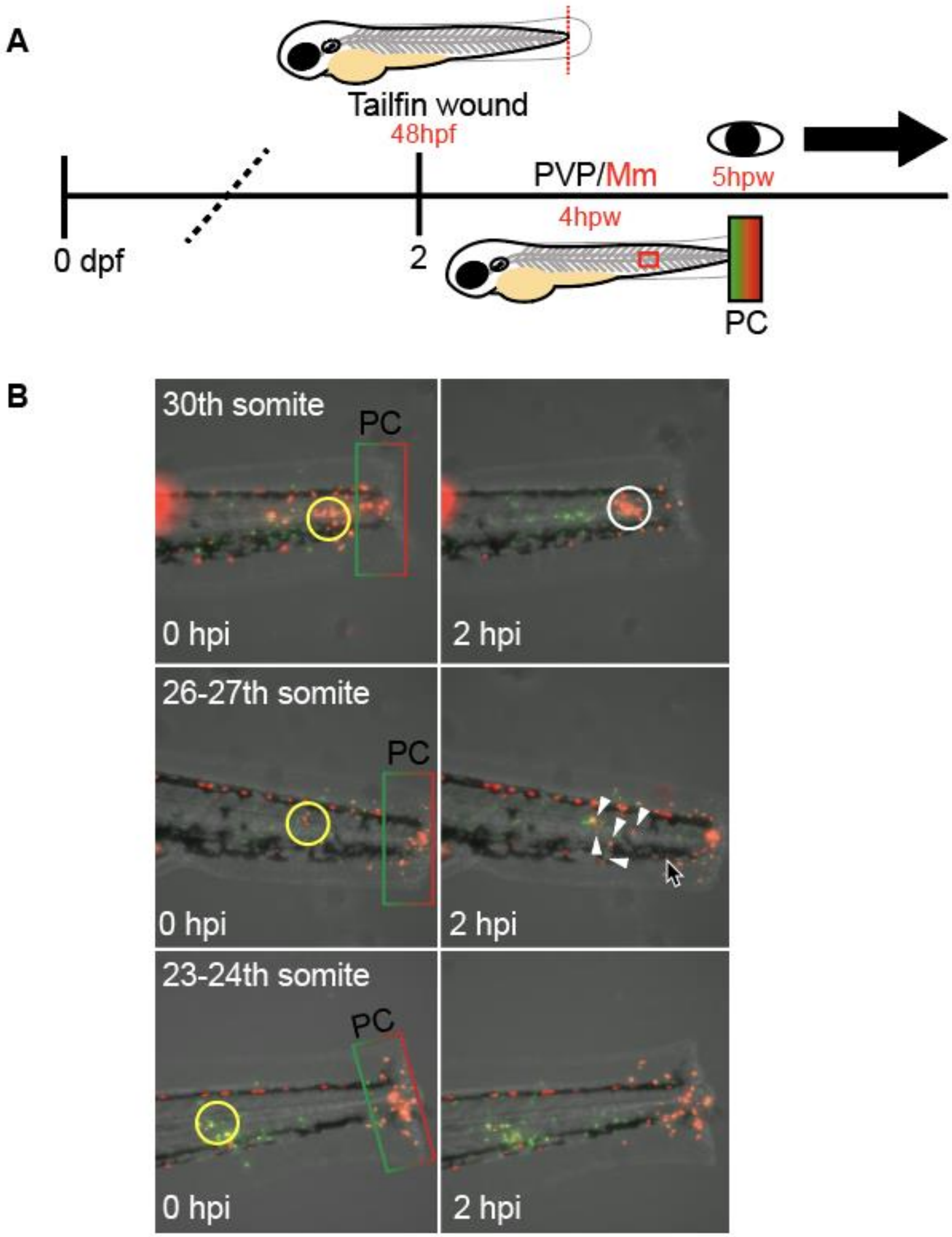

**Figure S1: Local Mm injection into the 26-27th somite attracted wound-experienced neutrophils away from a tailfin wound.**

(A) Schematic of experiment for B.

(B) Stereo-fluorescence micrographs of a tailfin transected embryo after either 30th somite, 26/27th somite, or 23-24 somite infection with Mm at 0hpi and 2hpi. The local infection site is shown by a yellow ring and photoconversion (PC) of wound neutrophils is shown by the box. With 30 somite injection red (wound-experienced) neutrophil have started migration to the infection site by the time the timelapse has been started and are almost all at the infection site by 2hpi (white ring). The 26th-27th somite infection does not start recruiting wound-experienced neutrophil by the time of the timelapse, but has done so after 2hpi (white arrowheads). Infection into the 23rd-24th somite does not recruit any wound-experienced neutrophils over a 2 hour time-course.

**Movie S1: A Hif-1$\alpha$ stabilised wound-experienced neutrophil with shuttling behaviour between the infection and wound sites.**

An example widefield fluorescence microscopy timelapse of a larvae with dominant active Hif-1$\alpha$, in which a red, wound-experienced neutrophil can be seen repeatedly migrating to and from the somite Mm infection site in a shuttling movement, a behaviour not observed in green, wound-naïve neutrophils.
